# Comparison of antihypertensive drug classes for dementia prevention

**DOI:** 10.1101/517482

**Authors:** Venexia M Walker, Neil M Davies, Richard M Martin, Patrick G Kehoe

## Abstract

**Introduction:** There is evidence that hypertension in midlife can increase the risk of Alzheimer’s disease and vascular dementia in late life. In addition, some treatments for hypertension have been proposed to have cognitive benefits, independent of their effect on hypertension. Consequently, there is potential to repurpose treatments for hypertension for dementia. This study systematically compared seven antihypertensive drug classes for this purpose, using data on over 849,000 patients from the Clinical Practice Research Datalink.

**Methods:** Treatments for hypertension were assessed in an instrumental variable (IV) analysis to address potential confounding and reverse causation. Physicians’ prescribing preference was used as a categorical instrument, defined by the physicians’ last seven prescriptions. Participants were new antihypertensive users between 1996-2016, aged 40 and over.

**Findings:** We analysed 849,378 patients with total follow up of 5,497,266 patient-years. Beta-adrenoceptor blockers and vasodilator antihypertensives were found to confer small protective effects – for example, vasodilator antihypertensives resulted in 27 (95% CI: 17 to 38; p=4.4e-7) fewer cases of any dementia per 1000 treated compared with diuretics.

**Interpretation:** We found small differences in antihypertensive drug class effects on risk of dementia outcomes. However, we show the magnitude of the differences between drug classes is smaller than previously reported. Future research should look to implement other causal analysis methods to address biases in conventional observational research with the ultimate aim of triangulating the evidence concerning this hypothesis.

**Funding:** This work was supported by the Perros Trust and the Integrative Epidemiology Unit. The Integrative Epidemiology Unit is supported by the Medical Research Council and the University of Bristol [grant number MC_UU_00011/1, MC_UU_00011/3].

**RESEARCH IN CONTEXT:** *Evidence before this study:* A recent systematic review and meta-analysis has collated the evidence for treating hypertension to prevent dementia. Seven comparable observational studies were identified that used either case-control designs with logistic regression or cohort designs with survival analysis. These studies suggested that some classes, such as angiotensin-II receptor blockers, may prevent dementia. However, conventional observational analyses, such as these, can be subject to confounding and reverse causation.

*Added value of this study:* We have provided new evidence about the potential effects of antihypertensives on risk of dementia through the novel application of instrumental variable analysis to this research question and have shown that the magnitude of the differences between drug classes is smaller than many observational studies have previously reported.

*Implications of all the available evidence:* Further research is needed to triangulate this evidence with other sources and to understand the inconsistencies between the studies conducted to date. Ultimately, this will inform the prioritization of antihypertensive drug classes for dementia prevention.

## INTRODUCTION

There is a substantial unmet clinical need for treatments for dementia where benefits to patients, society and the public purse can be gained. Despite this, some drug companies have recently withdrawn from this therapy area due to failed and costly efforts to find new treatments. (1) Drug repurposing, the identification of properties in existing or abandoned compounds for other clinical conditions, offers significant advantages over traditional drug discovery approaches. This includes immediate access to human safety data from the original clinical development work, which can accelerate testing in clinical trials, saving both time and money. (2,3)

Many antihypertensive medications have been proposed as drug repurposing candidates for the prevention of dementia. In part, because of research to better understand the observed associations between midlife hypertension and later-life risk of Alzheimer’s disease and vascular dementia. (3–6) There is also increasing recognition that one of the earliest pathological events in the development of Alzheimer’s disease is vascular dysregulation. [8] As well as suggestions that some antihypertensives, specifically those that block angiotensin receptor and calcium channel signalling, may have other neurological benefits. (7–9)

Several observational studies have investigated repurposing antihypertensives for dementia prevention. (10–17) However, these studies have used case-control designs with logistic regression and cohort designs with survival analysis, which may be subject to unmeasured or residual confounding and reverse causation. Specifically, confounding by indication, where the reasons that a patient receives a treatment relate to the reasons that the patient is at an increased risk of the outcome; and healthy adherer bias, where patients initiating or adhering to a drug for prevention of a condition are more likely to be healthy; are of concern in this type of study. There is also potential for reverse causation due to preclinical disease, which could raise blood pressure prior to clinical symptoms and consequently lead to the prescription of an antihypertensive drug.

Instrumental variable analysis, which estimates the causal effect of an exposure on an outcome by using a third variable (the instrument), can be robust to confounding and reverse causation if certain assumptions are met. That is, the instrument must: (i) be associated with the exposure of interest; (ii) affect the outcome only through its effect on the exposure; and (iii) have no common causes with the outcome (i.e. there are no confounders of the instrument-outcome association). (18,19) Physicians’ prescribing preference has been proposed as an instrumental variable in pharmacoepidemiology. (20–24) It meets the instrument conditions as: (i) it is associated with the prescription issued by the physician; (ii) it is unlikely to relate to the patient’s risk of dementia other than through the prescription issued; and (iii) physicians’ prescribing preference is unlikely to share a cause with the patient’s outcome because patients have relatively little choice over which physician they see or knowledge of their physicians’ preferences for antihypertensive drug classes. (22) We therefore report a systematic assessment of the major antihypertensive drug classes as candidates for the prevention of Alzheimer’s disease, vascular dementia, other dementias and any dementia, using physicians’ prescribing preference as an instrument in the Clinical Practice Research Datalink (CPRD) to overcome confounding and reverse causation.

## METHODS

### Study design

We conducted a prospective new user cohort study in the CPRD. (25) The CPRD is a primary care database with over 11.3 million people from more than 670 UK practices. (26) The data were extracted from the CPRD-GOLD primary care dataset March 2016 snapshot (ISAC 15_246R). This snapshot included all patients with ‘research quality’ data, who registered at a participating practice from 1^st^ January 1987 to 29^th^ February 2016. (27) The *a priori* protocol for this study was published prior to the current report (see Supplementary Table 1 for amendments) and the study design diagram is included as Supplementary Figure 1. (28)

### Participants

Patients were included in the analysis if they were aged 40 years or over and received a first prescription for an antihypertensive drug class of interest. Follow-up was stopped at the earliest of: a dementia outcome; death; end of registration at a CPRD general practice; or the end of follow-up for this study (29^th^ February 2016). Patients were excluded if they were of unknown gender; had less than 12 months of ‘research quality’ data prior to their first prescription (to improve the identification of baseline covariates); or were initially prescribed multiple antihypertensive drug class of interest. We also excluded patients prescribed an antihypertensive before 1^st^ January 1996, as 1996 was the first complete year that all of the drugs being considered were available.

### Exposures

We considered seven antihypertensive drug classes based on the groupings in the British National Formulary. (29) These were: alpha-adrenoceptor blockers, angiotensin-converting enzyme inhibitors, angiotensin-II receptor blockers, beta-adrenoceptor blockers, calcium channel blockers, diuretics (either ‘thiazides and related diuretics’ or ‘potassium-sparing diuretics and aldosterone antagonists’), and vasodilator antihypertensives. To mimic a randomised controlled trial (RCT), exposure to the drug classes was analysed in an intention-to-treat framework, i.e. based on the first prescription irrespective of subsequent switches to, or additions of, other antihypertensive drug classes. (30) The index date for each patient was the date they received their first prescription for an antihypertensive drug. Treatment switching was not modelled, as it was likely to be non-random and confounded by patients’ unobservable characteristics.

### Outcomes

We defined four outcomes for this analysis: probable Alzheimer’s disease, possible Alzheimer’s disease, vascular dementia and other dementias (Supplementary Figure 2). We also considered any dementia, which combined the dementia subtypes in a single outcome.

### Covariates

The instrumental variable analysis was adjusted for prescription year only. This was necessary as the number of antihypertensive prescriptions in the CPRD varied by year and so may have influenced both the instrument-exposure and instrument-outcome associations. All other potential covariates were thought to influence the exposure-outcome association, but not the instrument-exposure or instrument-outcome associations, and so will be balanced across levels of the instrument if the instrument assumptions are met. The instrumental variable analysis was compared with a multivariable logistic regression analysis to assess the extent of confounding. The multivariable logistic regression analysis was adjusted for prescription year; sex; age at index; previous history of coronary heart disease, coronary-bypass surgery, or cerebrovascular disease; chronic disease; socioeconomic position; consultation rate; alcohol status; smoking status; and body mass index (BMI). All covariates were determined prior to index and are defined fully in Supplementary Table 3.

### Code lists

Prescriptions and diagnoses were defined using Product and Read codes respectively. These codes are recorded at the time of the consultation and uniquely define prescriptions and clinical terms in the CPRD. The code lists for this study are provided on Github (https://github.com/venexia/repurposing-antihypertensives-dementia).

### Assessment of bias

To assess bias, we constructed bias scatter plots for each outcome. These plots compare the association of each covariate with the exposure (obtained from multivariable linear regression analysis) and the instrument (obtained from instrumental variable analysis). (31,32) See Supplementary Text 1 for interpretation. Any covariates found to be as, or more, biased for the instrumental variable analysis (i.e. on or above the x=y line) were adjusted for in a sensitivity analysis.

### Statistical methods

This study used instrumental variable analysis with physicians preferred antihypertensive drug class as an instrument to proxy for exposure, i.e. the actual drug class prescribed (Figure 1). Each drug class was used as the reference drug class for each of the other drug classes in a series of pairwise comparisons. Prescribing preference was derived from the prescriptions issued by the physician to their seven most recent patients who received an antihypertensive. (33,34) This resulted in an ordered categorical instrument indicating how many previous prescriptions the physician had issued for the drug class of interest over the reference drug class in the present pairwise comparison. The analysis used the ivreg2 package in Stata with ‘robust’ specified (to address arbitrary heteroskedasticity) and clustering by physician (to address both arbitrary heteroskedasticity and intra-group correlations between physicians). (35) To obtain a point estimate, we made a further assumption - in addition to the three standard instrument assumptions - of monotonicity. That is, we assumed all patients complied with their physicians’ preferred drug class. Consequently, the results were interpreted as the effect among patients whose prescription was affected by their physicians’ preference (known as the local average treatment effect). For each analysis, we present the partial F statistic to quantify and test the strength of the instrument-exposure association. We also present the results of endogeneity tests conducted using the option ‘endog’ in ivreg2. The analysis is presented in line with reporting guidelines (Supplementary Table 2). (36) All analyses were conducted in Stata version 15MP and R version 3.4.4. (37,38) The code is available from GitHub (https://github.com/venexia/repurposing-antihypertensives-dementia).

**Figure 1:**
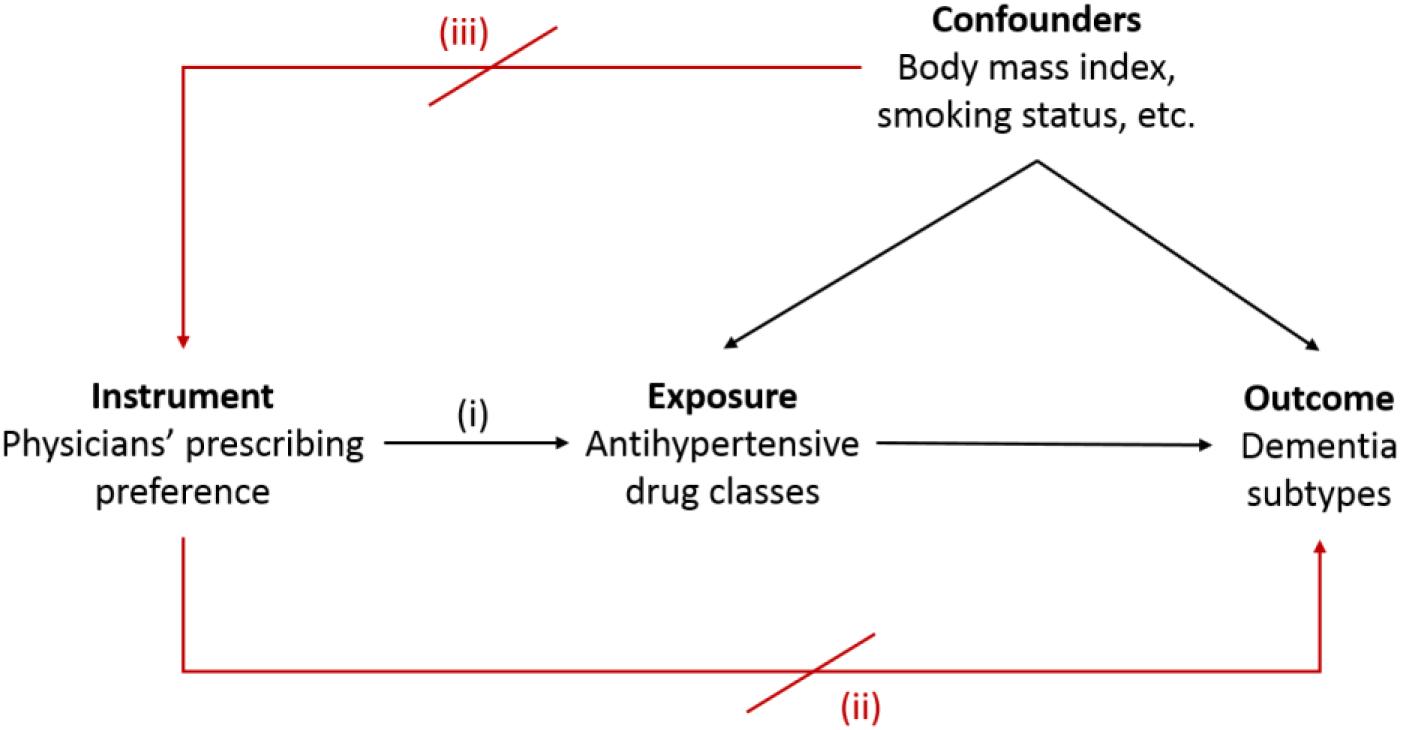
Directed acyclic graph for the instrumental variable analysis. Instrumental variable analysis requires that the instrument: (i) be associated with the exposure of interest; (ii) affect the outcome only through its effect on the exposure of interest; and (iii) have no common causes with the outcome. To obtain a point estimate for this analysis, we also make a fourth assumption of monotonicity. The measured confounders of this analysis are listed in the section ‘covariates’, however there is also likely to be unmeasured confounders of the exposure-outcome association hence warranting the use of this method.

### Sensitivity analyses

Beta-adrenoceptor blockers can be prescribed in low doses for the treatment of anxiety. (39). However, to be a suitable comparator, we required them to be prescribed for the treatment of hypertension. We therefore tested the effect of removing patients thought to be receiving these drugs for anxiety in two ways. Firstly, we did the analysis without people who both received a drug class of interest and had a Read code indicating anxiety, or other neurotic, stress-related and somatoform disorders in the same consultation (using a previously published code list). (40) Secondly, we did the analysis without people whose dose was in the bottom 25% for their index drug class.

Differential prescribing occurs in women of child bearing age due to risks associated with some antihypertensives during pregnancy. (41) As participants can enter this study at the age of 40, this might affect the youngest members of the cohort. We therefore conducted a sensitivity analysis restricted to patients aged 55 and over at index. This age threshold is currently being used in the RADAR trial for similar reasons. (42)

## RESULTS

### Patient characteristics

A total of 849,378 patients, with a total follow up of 5,497,266 patient years, met the criteria for our analysis. Supplementary Figure 3 outlines patient attrition. Table 1 presents patient characteristics. (43–45) The full cohort had a median age of 61 (interquartile range: 51-71) at index date and a median follow-up of 5.8 years (interquartile range: 2.6-9.8). Of the 849,378 patients, 410,805 (48%) had complete covariate information. This subset of patients was used when comparing instrumental variable and multivariable logistic regression analyses. The subset had a median age of 61 (interquartile range: 51-71) at index date, and median follow-up of 5.6 years (interquartile range: 2.5-9.5). Incomplete covariate information was mainly due to missing values for the Index of Multiple Deprivation (IMD), which was used to adjust for socioeconomic position, as this measure is only available for patients in English practices. One notable feature of the patient characteristics is that 97% of patients receiving alpha-adrenoceptor blockers and 99.5% of patients receiving vasodilator antihypertensives were men - this difference persists regardless of the age at first prescription (Supplementary Table 4).

**Table 1:** Patient characteristics

|  | Alpha-adrenoceptor blockers | Angiotensin-II receptor blockers | Angiotensin-converting enzyme inhibitors | Beta-adrenoceptor blockers | Calcium channel blockers | Diuretics | Vasodilator anti-hypertensives | Whole sample |
| --- | --- | --- | --- | --- | --- | --- | --- | --- |
| N | 67360 | 14717 | 195891 | 240864 | 139730 | 180946 | 9870 | 849378 |
| Median year of first prescription | 2008 | 2005 | 2007 | 2005 | 2008 | 2003 | 2008 | 2006 |
| Male sex | 97.0%<br>(65365) | 55.3%<br>(8141) | 58.0%<br>(113667) | 43.2%<br>(104096) | 49.2%<br>(68739) | 36.0%<br>(65177) | 99.3%<br>(9796) | 51.2%<br>(434981) |
| Median age at first prescription | 65 | 59 | 59 | 55 | 64 | 66 | 57 | 61 |
| Previous history of coronary artery disease | 0.2%<br>(129) | 0.6%<br>(85) | 0.8%<br>(1536) | 0.9%<br>(2056) | 0.4%<br>(562) | 0.1%<br>(203) | 0.1%<br>(11) | 0.5%<br>(4582) |
| Previous history of coronary-bypass surgery | 0.3%<br>(193) | 0.3%<br>(45) | 0.5%<br>(946) | 0.5%<br>(1262) | 0.3%<br>(418) | 0.1%<br>(265) | 0.1%<br>(14) | 0.4%<br>(3143) |
| Previous history of cerebrovascular disease | 2.0%<br>(1319) | 2.1%<br>(311) | 3.0%<br>(5813) | 1.4%<br>(3387) | 2.3%<br>(3194) | 2.8%<br>(5090) | 0.7%<br>(73) | 2.3%<br>(19187) |
| At least one comorbidity on the Charlson index<sup>a</sup> | 36.8%<br>(24817) | 42.4%<br>(6238) | 50.8%<br>(99492) | 26.0%<br>(62604) | 38.7%<br>(54081) | 36.0%<br>(65212) | 42.6%<br>(4207) | 37.3%<br>(316651) |
| Median IMD 2010 score<sup>b</sup> | 8 | 8 | 9 | 9 | 9 | 9 | 8 | 9 |
| Mean annual consultation rate (SD) | 5.6<br>(5.4) | 6.1<br>(6.3) | 6.1<br>(6.0) | 5.8<br>(5.3) | 5.9<br>(5.8) | 6.0<br>(5.6) | 5.5<br>(5.1) | 5.9<br>(5.7) |
| Ever drinker<sup>c</sup> | 89.2%<br>(60070) | 85.2%<br>(12538) | 85.6%<br>(167636) | 86.1%<br>(207457) | 84.5%<br>(118104) | 84.3%<br>(152473) | 91.8%<br>(9059) | 85.6%<br>(727337) |
| Ever smoker<sup>d</sup> | 54.5%<br>(36691) | 52.5%<br>(7729) | 53.8%<br>(105401) | 54.3%<br>(130894) | 53.3%<br>(74540) | 55.2%<br>(99793) | 57.6%<br>(5688) | 54.2%<br>(460736) |
| Mean BMI (SD)<sup>e</sup> | 26.5<br>(4.2) | 28.6<br>(5.7) | 29.0<br>(5.9) | 26.6<br>(5.0) | 27.5<br>(5.4) | 27.5<br>(5.5) | 27.3<br>(4.4) | 27.5<br>(5.4) |

- (a) The Charlson index is a classification of 17 chronic diseases, including cancer and arthritis, which may alter mortality risk. - (b) IMD 2010 score is a proxy for socioeconomic position that is measured as 'twentiles' with 1 indicating the least deprived and 20 indicating the most deprived. IMD 2010 score was missing for 38.6% (328,233) of the whole sample. - (c) Alcohol status was missing for 15.6% (132,387) of the whole sample. For the purposes of this table, it has been classified as 'ever' (i.e. former or current) vs 'never'. - (d) Smoking status was missing for 6.4% (54,447) of the whole sample. For the purposes of this table, it has been classified as 'ever' (i.e. former or current) vs 'never'. - (e) BMI, or a calculated BMI from height and weight measurements, was missing for 15.7% (128,830) of the whole sample.

### Alzheimer’s disease

Figure 2 shows the results for probable and possible Alzheimer’s disease respectively. Our results suggested that beta-adrenoceptor blockers were protective for both probable and possible Alzheimer’s disease when compared with other drugs. For example, beta-adrenoceptor blockers were estimated to result in 8 (95% CI: 3 to 12; p=3.1e-3) fewer cases of probable Alzheimer’s disease and 9 (95% CI: 4 to 13; p=1.3e-4) fewer cases of possible Alzheimer’s disease per 1000 people treated when compared with alpha-adrenoceptor blockers.

**Figure 2:**
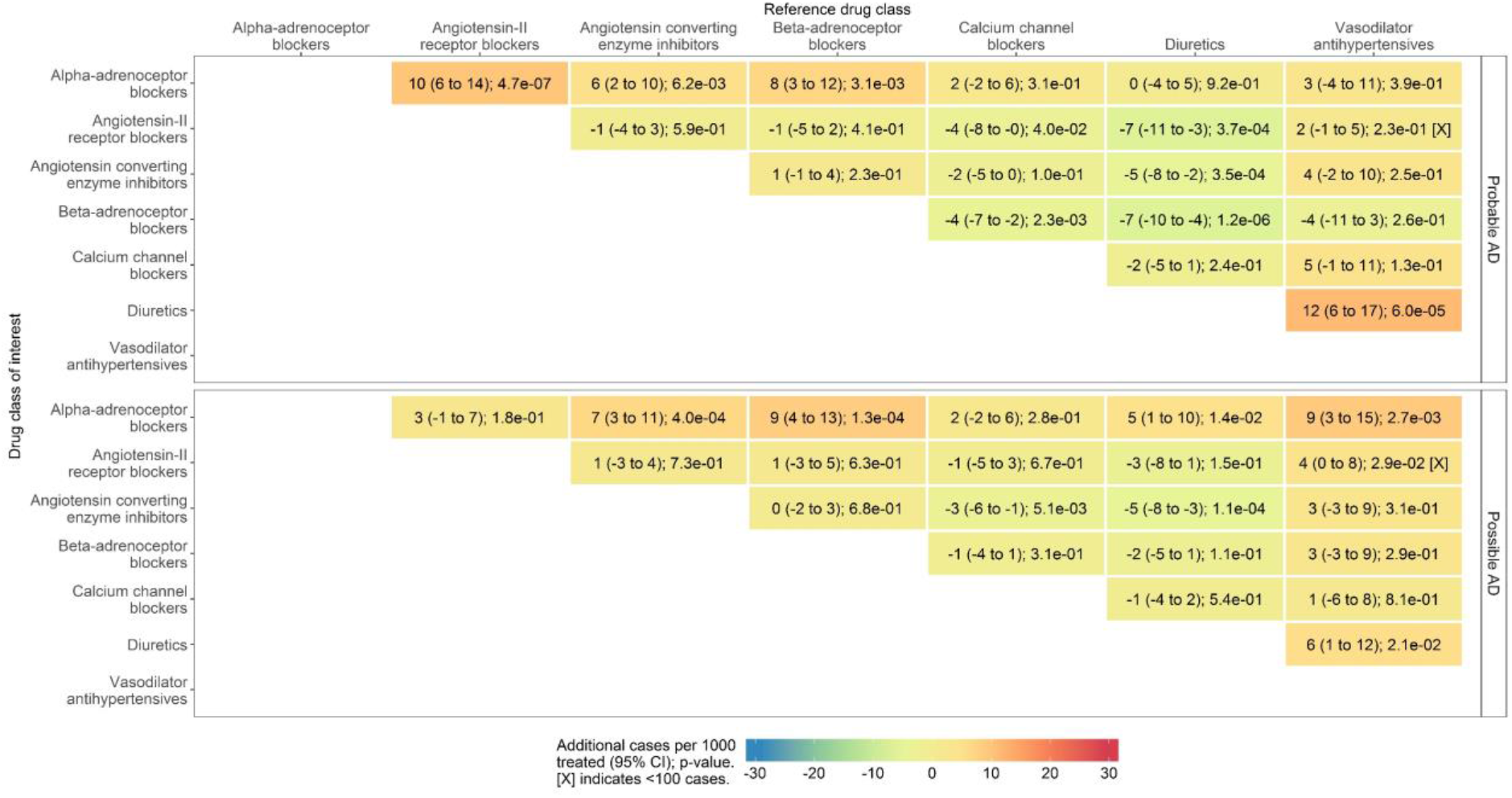
Instrumental variable estimates for the risk of probable and possible Alzheimer’s disease. F greater than 4708 for all analyses (Supplementary Table 5).

### Non-Alzheimer’s disease dementias

Figure 3 shows the results for vascular and other dementias respectively. The magnitude of the differences between drug classes is smaller for these outcomes. However, vasodilator antihypertensives were suggested to be protective with an estimated 5 (95% CI: 0 to 9; p=4.0e-2) fewer cases of vascular dementia and 6 (95% CI: 1 to 11; p=1.5e-2) fewer cases of other dementias per 1000 people treated when compared with calcium channel blockers. Angiotensin-II receptor blockers were also indicated to be protective for vascular dementia with an estimated 7 (95% CI: 4 to 10; p=1.4e-5) fewer cases of vascular dementia per 1000 people treated when compared with alpha-adrenoceptor blockers.

**Figure 3:**
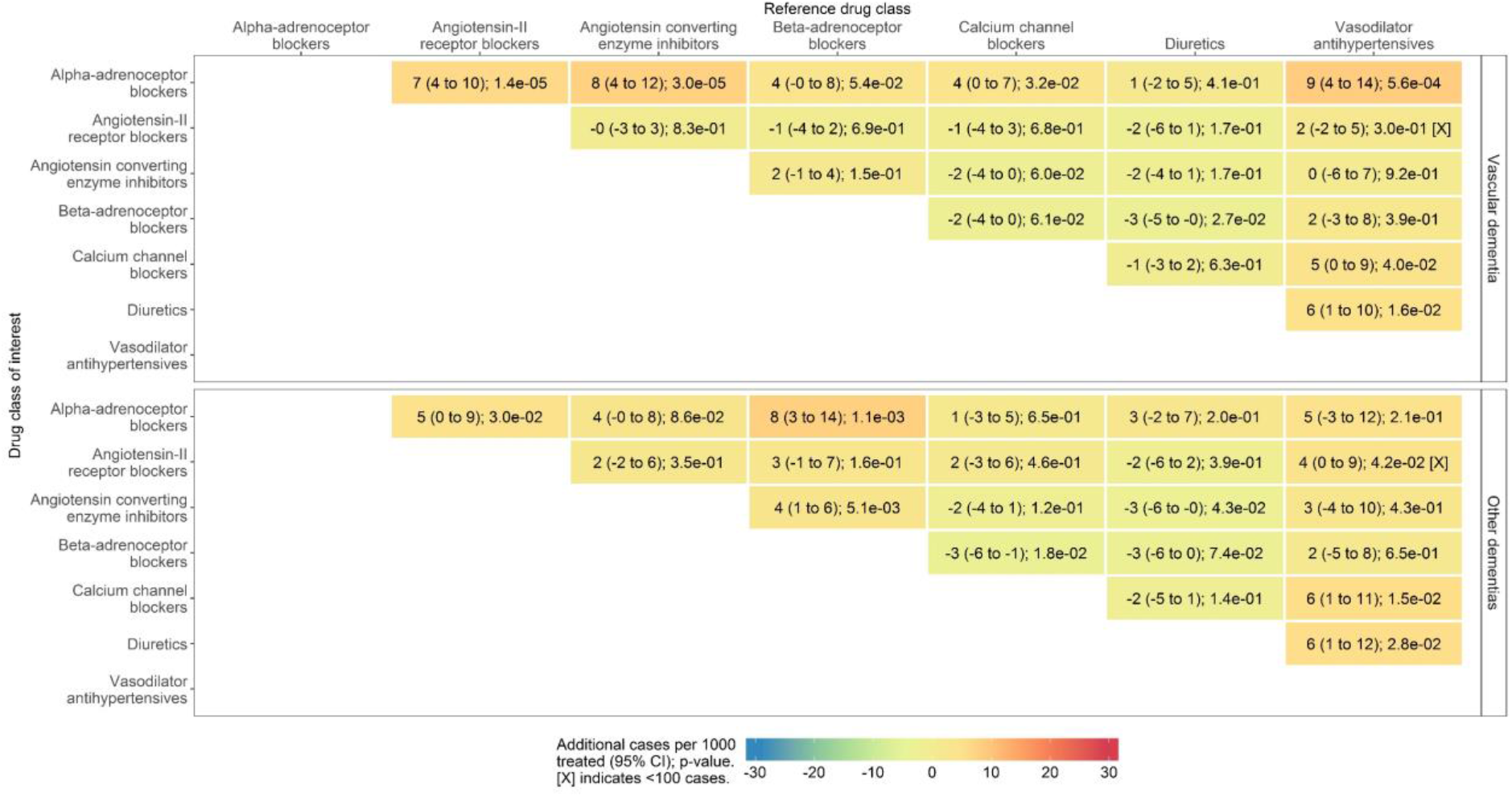
Instrumental variable estimates for the risk of non-Alzheimer’s disease dementia. F greater than 4702 for all analyses (Supplementary Table 5).

### Any dementia

Figure 4 shows the results for any dementia. These results reflected the dementia subtype analyses and emphasised the effects observed, perhaps due to the increased sample size. For example, beta-adrenoceptor blockers were estimated to result in 28 (95% CI: 19 to 38; p=5.2e-9) fewer cases per 1000 people treated compared with alpha-adrenoceptor blockers. Meanwhile, vasodilator antihypertensives were estimated to result in 27 (95% CI: 17 to 38; p=4.4e-7) fewer cases per 1000 people treated compared with diuretics.

**Figure 4:**
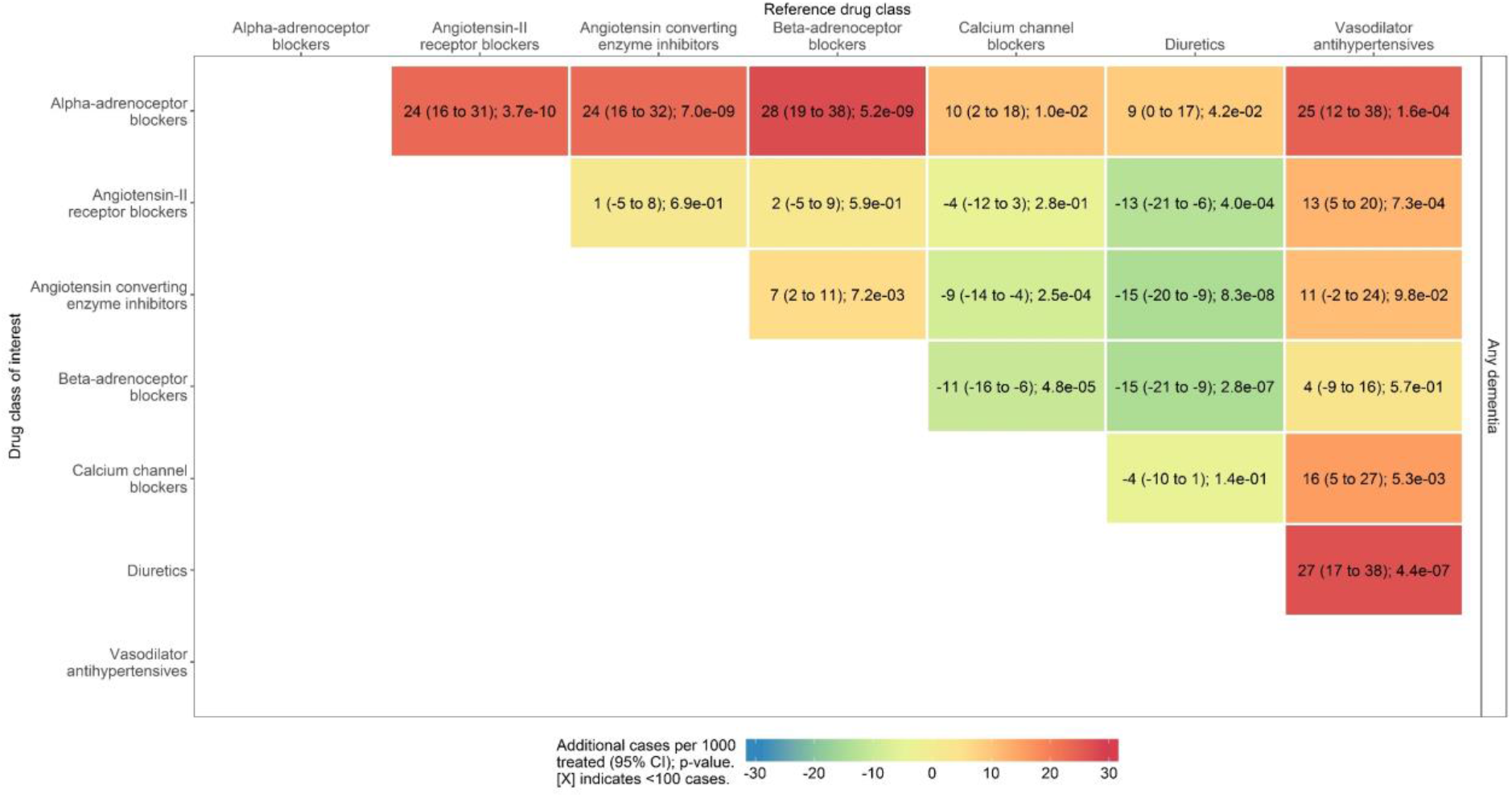
instrumental variable estimates for the risk of any dementia. F greater than 4876 for all analyses (Supplementary Table 5).

### Comparison with multivariable logistic regression

The results of the multivariable logistic regression are provided in Supplementary Figure 4. Endogeneity tests indicated evidence to reject the null that the exposure was endogenous, indicating a difference between the instrumental variable analysis and ordinary least squares results, for a small number of the analyses run (Supplementary Table 5). Most of these analyses considered alpha-adrenoceptor blockers as the drug class of interest.

### Assessment of bias

Bias scatter plots were used to assess bias among the subset of patients with complete covariate information (Supplementary Figure 5). The bias term was larger in the instrumental variable analysis, compared to the multivariable linear regression analysis, for socioeconomic position only. Bias terms were equally biased for BMI, chronic disease, sex and age. These covariates, including socioeconomic position, were adjusted for in sensitivity analyses and were mostly found to produce consistent results with the main analysis (Supplementary Figures 6-10). The exception was results concerning diuretics and beta-adrenoceptor blockers after adjustment for age. These drug classes have the oldest and youngest median ages at index respectively (Table 1), which may explain why they were most effected by the adjustment.

### Sensitivity analyses

There was minimal effect of removing those diagnosed with anxiety in the same consultation from our analysis (Supplementary Figure 11). Similarly, we observed little difference after removing those who received a low dose initial prescription though there was a lack of power for some analyses (Supplementary Figure 12). Finally, restricting the analysis to patients aged 55 and over at index did not change the direction of effect for our results however, several effects failed to exclude the null after being subject to this restriction (Supplementary Figure 13).

## DISCUSSION

### Principal findings

Beta-adrenoceptor blockers and vasodilator antihypertensives reduced risk of probable and possible Alzheimer’s disease, vascular dementia, other dementias, and any dementia when compared with other antihypertensive drug classes. On the contrary, diuretics and alpha-adrenoceptor blockers increased risk of dementia outcomes when compared with other antihypertensive drug classes. Our results concerning beta-adrenoceptor blockers and diuretics may be biased by age, however this bias is no more extreme than that observed for multivariable linear regression. This study does not explore the effect of antihypertensives treatment compared to non-treatment on risk of dementia, which a meta-analysis of RCTs suggests has a relative risk of 0.84 (95% CI: 0.69 to 1.02; p=0.10). (10)

### Comparison with existing literature

There is currently one published RCT with Alzheimer’s disease as a primary outcome that has assessed whether an antihypertensive drug, nilvadipine, could benefit patients. This trial compared the treatment against non-use and failed to show treatment benefit. (46) There have been no RCTs published to date that have directly compared antihypertensive drug classes to each other for the prevention or treatment of Alzheimer’s disease. However, a recent meta-analysis by Larsson et al identified seven prospective observational studies on this topic. (10–17) Two of which also made use of the CPRD (Supplementary Text 2). (13,16) Relative to other antihypertensive drug classes: one study (of three) suggested angiotensin-converting enzyme inhibitors were protective (11–13); three studies (of four) suggested angiotensin-II receptor blockers were protective (13–16); and one study (of one) suggested calcium channel blockers were protective (17). In contrast, our analysis suggested beta-adrenoceptor blocker and vasodilator antihypertensives were among the most protective drug classes when compared with other antihypertensives. Since this meta-analysis was published, Barthold et al have conducted a study comparing Alzheimer’s disease incidence between users of renin-angiotensin system (RAS) acting drug classes (angiotensin-converting enzyme inhibitors and angiotensin-II receptor blockers) and non-RAS acting drug classes (beta blockers, calcium channel blockers, loop diuretics, and thiazide-like diuretics) across sex, race, and ethnic groups in the United States. (47) They found that angiotensin-II receptor blockers may reduce the risk of Alzheimer’s disease in certain groups, namely white and black women and white men. Our study is in a population of mainly white men and women and did not find such a clear distinction between angiotensin-II receptor blockers and non-RAS acting drugs. As already highlighted, the major difference between our observational study and those previously conducted is the statistical methods used. When the analysis assumptions are met, our instrumental variable analysis should not be subject to unmeasured confounding, which may affect other types of analysis.

### Strengths and limitations

The key strength of this study was the large cohort of patients (consisting of 849,378 patients with 5,497,266 patient years of follow-up) that would not be achievable in RCTs.

The size of our study meant we had ample power to detect even small differences between the drug classes of interest. We also had data on both male and female patients unlike some of the larger existing studies. (14) In addition, we used instrumental variable analysis, which should not be subject to unmeasured confounding when the assumptions hold, and an active comparator design, whereby antihypertensive drug classes were compared with other antihypertensive drug classes, to ensure patients were comparable. (25,30) These analysis features were important in the present study due to the risk of confounding.

Despite finding some evidence of bias, the sensitivity analyses showed the effect on our results to be minimal with only minor changes in magnitude for most estimates. This includes the potential bias due to socioeconomic position, which was deemed the most extreme in our assessment. The only concern was potential confounding by age for results relating to beta-adrenoceptor blockers and diuretics, however this bias was no more extreme than that observed in the multivariable linear regression.

A limitation of our study is we cannot prove that the instrumental variable assumptions hold. The only assumption that can be empirically tested is the first, namely that the instrument is associated with the rates of prescribing. Our proposed instruments had a minimum F statistic of 4702 in our main analyses, demonstrating they strongly associated with the exposure. Our study may also have misclassified the exposure due to the use of the intention-to-treat framework, which defines exposure based on the first treatment prescribed. However, the benefits of this approach – such as preserving sample size and replicating ‘real world’ prescribing – outweigh the concerns. Finally, this study may have misclassified outcomes, which can occur when a diagnosis is not updated or recorded accurately in primary care records. We took steps to overcome this by considering ‘probable’ and ‘possible’ definitions for Alzheimer’s disease, the most common form of dementia. We also included an ‘any dementia’ outcome that should not be affected by the difficulties of determining subtype.

### Conclusions and implications

We have provided new evidence about the potential effects of antihypertensives on risk of dementia through the novel application of instrumental variable analysis to this research question. We found small differences in drug class effects on risk of dementia outcomes. For example, we found that beta-adrenoceptor blockers were estimated to result in 15 (95% CI: 9 to 21; p=2.8e-7) fewer cases of any dementia per 1000 people treated compared with diuretics and 11 (95% CI: 6 to 16; p=4.8e-5) fewer when compared with calcium channel blockers. However, we show the magnitude of the differences between drug classes is smaller than many observational studies have previously reported. Future research should identify potential sources of unmeasured confounding that may have affected previous observational studies to understand this inconsistency. This may also provide a stimulus for more in-depth investigations of the related biological mechanisms, which will in turn inform the study of both the disease process and potential drug targets for dementia prevention.

## Supporting information

Codelists

## AUTHORS’ CONTRIBUTIONS

All authors contributed to planning the analysis. VMW conducted the analysis and drafted the manuscript. All other authors edited and revised the manuscript. PGK and RMM were responsible for securing the funding. All authors read and approved the final manuscript.

## ACKNOWLEDGEMENTS

This study is based in part on data from the Clinical Practice Research Datalink obtained under licence from the UK Medicines and Healthcare products Regulatory Agency. However, the interpretation and conclusions contained in this report are those of the authors alone.

## REFERENCES

1. Jobke B, McBride T, Nevin L, Peiperl L, Ross A, Stone C, et al. Setbacks in Alzheimer research demand new strategies, not surrender. PLOS Med. 2018 Feb 27;15(2):e1002518.

2. S. B Appleby, L. J. Cummings Discovering New Treatments for Alzheimer’s Disease by Repurposing Approved Medications. Curr Top Med Chem. 2013 Sep 1;13(18):2306–27.

3. Corbett A, Pickett J, Burns A, Corcoran J, Dunnett SB, Edison P, et al. Drug repositioning for Alzheimer’s disease. Nat Rev Drug Discov. 2012;11(11):833–846.

4. Abell JG, Kivimäki M, Dugravot A, Tabak AG, Fayosse A, Shipley M, et al. Association between systolic blood pressure and dementia in the Whitehall II cohort study: role of age, duration, and threshold used to define hypertension. Eur Heart J [Internet]. [cited 2018 Jun 13]; Available from: https://academic.oup.com/eurheartj/advance-article/doi/10.1093/eurheartj/ehy288/5032485

5. Kennelly SP, Lawlor BA, Kenny RA. Blood pressure and the risk for dementia—A double edged sword. Ageing Res Rev. 2009 Apr 1;8(2):61–70.

6. Qiu C, Winblad B, Fratiglioni L. The age-dependent relation of blood pressure to cognitive function and dementia. Lancet Neurol. 2005 Aug;4(8):487–99.

7. Kehoe PG, Passmore PA. The renin-angiotensin system and antihypertensive drugs in Alzheimer’s disease: current standing of the angiotensin hypothesis? J Alzheimers Dis JAD. 2012;30 Suppl 2:S251–268.

8. Kehoe PG, Miners S, Love S. Angiotensins in Alzheimer’s disease - friend or foe? Trends Neurosci. 2009 Dec;32(12):619–28.

9. Wright AK, Kontopantelis E, Emsley R, Buchan I, Sattar N, Rutter MK, et al. Life Expectancy and Cause-Specific Mortality in Type 2 Diabetes: A Population-Based Cohort Study Quantifying Relationships in Ethnic Subgroups. Diabetes Care. 2017 Mar 1;40(3):338–45.

10. Larsson SC, Markus HS. Does Treating Vascular Risk Factors Prevent Dementia and Alzheimer’s Disease? A Systematic Review and Meta-Analysis. J Alzheimers Dis. 2018 Jan 1;64(2):657–68.

11. Ohrui T, Matsui T, Yamaya M, Arai H, Ebihara S, Maruyama M, et al. Angiotensin-converting enzyme inhibitors and incidence of Alzheimer’s disease in Japan. J Am Geriatr Soc. 2004 Apr;52(4):649–50.

12. Sink KM, Leng X, Williamson J, Kritchevsky SB, Yaffe K, Kuller L, et al. Angiotensin-converting enzyme inhibitors and cognitive decline in older adults with hypertension: results from the Cardiovascular Health Study. Arch Intern Med. 2009 Jul 13;169(13):1195–202.

13. Davies NM, Kehoe PG, Ben-Shlomo Y, Martin RM. Associations of anti-hypertensive treatments with Alzheimer’s disease, vascular dementia, and other dementias. J Alzheimers Dis. 2011;26(4):699.

14. Li N, Lee A, Whitmer R, Kivipelto M, Lawler E, Kazis L, et al. Use of angiotensin receptor blockers and risk of dementia in a predominantly male population: prospective cohort analysis. Br Med J. 2010;340(jan12 1):b5465.

15. Hsu C-Y, Huang C-C, Chan W-L, Huang P-H, Chiang C-H, Chen T-J, et al. Angiotensin-receptor blockers and risk of Alzheimer’s disease in hypertension population--a nationwide cohort study. Circ J Off J Jpn Circ Soc. 2013;77(2):405–10.

16. Goh KL, Bhaskaran K, Minassian C, Evans SJW, Smeeth L, Douglas IJ. Angiotensin receptor blockers and risk of dementia: cohort study in UK Clinical Practice Research Datalink. Br J Clin Pharmacol. 2015 Feb 1;79(2):337–50.

17. Hwang D, Kim S, Choi H, Oh I-H, Kim BS, Choi HR, et al. Calcium-Channel Blockers and Dementia Risk in Older Adults - National Health Insurance Service - Senior Cohort (2002-2013). Circ J Off J Jpn Circ Soc. 2016 Oct 25;80(11):2336–42.

18. Hernán MA, Robins JM. Instruments for Causal Inference: An Epidemiologist’s Dream? Epidemiology. 2006 Jul;17(4):360–72.

19. Hernán M, Robins J. Causal Inference. Boca Raton: Chapman & Hall/CRC; forthcoming.

20. Schneeweiss S, Setoguchi S, Brookhart A, Dormuth C, Wang PS. Risk of death associated with the use of conventional versus atypical antipsychotic drugs among elderly patients. CMAJ Can Med Assoc J J Assoc Medicale Can. 2007 Feb 27;176(5):627–32.

21. Wang PS, Schneeweiss S, Avorn J, Fischer MA, Mogun H, Solomon DH, et al. Risk of death in elderly users of conventional vs. atypical antipsychotic medications. N Engl J Med. 2005 Dec 1;353(22):2335–41.

22. Davies NM, Gunnell D, Thomas KH, Metcalfe C, Windmeijer F, Martin RM. Physicians’ prescribing preferences were a potential instrument for patients’ actual prescriptions of antidepressants. J Clin Epidemiol. 2013 Dec;66(12):1386–96.

23. Schneeweiss S, Seeger JD, Landon J, Walker AM. Aprotinin during coronary-artery bypass grafting and risk of death. N Engl J Med. 2008 Feb 21;358(8):771–83.

24. Taylor GM, Taylor AE, Thomas KH, Jones T, Martin RM, Munafò MR, et al. The effectiveness of varenicline versus nicotine replacement therapy on long-term smoking cessation in primary care: a prospective cohort study of electronic medical records. Int J Epidemiol. 2017 Dec 1;46(6):1948–57.

25. Lund JL, Richardson DB, Stürmer T. The active comparator, new user study design in pharmacoepidemiology: historical foundations and contemporary application. Curr Epidemiol Rep. 2015 Dec;2(4):221–8.

26. Herrett E, Gallagher AM, Bhaskaran K, Forbes H, Mathur R, van Staa T, et al. Data Resource Profile: Clinical Practice Research Datalink (CPRD). Int J Epidemiol. 2015 Jun;44(3):827–36.

27. Walker V, Davies N, Kehoe P, Martin R. CPRD codes: neurodegenerative diseases and commonly prescribed drugs. 2017; Available from: https://doi.org/10.5523/bris.1plm8il42rmlo2a2fqwslwckm2

28. Walker VM, Davies NM, Jones T, Kehoe PG, Martin RM. Can commonly prescribed drugs be repurposed for the prevention or treatment of Alzheimer’s and other neurodegenerative diseases? Protocol for an observational cohort study in the UK Clinical Practice Research Datalink. BMJ Open. 2016 Dec 1;6(12):e012044.

29. BNF Legacy. BNF July 2017: BNF Legacy [Internet]. [cited 2018 Feb 16]. Available from: https://www.medicinescomplete.com/mc/bnflegacy/64/

30. Hernán MA, Robins JM. Using Big Data to Emulate a Target Trial When a Randomized Trial Is Not Available. Am J Epidemiol. 2016 Apr 15;183(8):758–64.

31. Jackson JW, Swanson SA. Toward a Clearer Portrayal of Confounding Bias in Instrumental Variable Applications: Epidemiology. 2015 Jul;26(4):498–504.

32. Davies NM, Thomas KH, Taylor AE, Taylor GM, Martin RM, Munafò MR, et al. How to compare instrumental variable and conventional regression analyses using negative controls and bias plots. Int J Epidemiol. 2017 Dec 1;46(6):2067–77.

33. Brookhart MA, Wang P, Solomon DH, Schneeweiss S. Evaluating short-term drug effects using a physician-specific prescribing preference as an instrumental variable. Epidemiol Camb Mass. 2006;17(3):268–75.

34. Brookhart MA, Schneeweiss S. Preference-based instrumental variable methods for the estimation of treatment effects: assessing validity and interpreting results. Int J Biostat. 2007;3(1):14.

35. Baum CF, Schaffer ME, Stillman S. ivreg2: Stata module for extended instrumental variables/2SLS, GMM and AC/HAC, LIML and k-class regression. [Internet]. 2010 [cited 2017 Nov 9]. Available from: http://ideas.repec.org/c/boc/bocode/s425401.html

36. Davies NM, Davey Smith G, Windmeijer F, Martin RM. Issues in the reporting and conduct of instrumental variable studies: a systematic review. Epidemiol Camb Mass. 2013 May;24(3):363–9.

37. StataCorp. Stata Statistical Software. College Station, TX: StataCorp LP; 2015.

38. R Core Team. R: A Language and Environment for Statistical Computing [Internet]. Vienna, Austria: R Foundation for Statistical Computing; Available from: https://www.R-project.org

39. The National Institute for Health and Care Excellence. Beta-adrenoceptor blocking drugs [Internet]. [cited 2018 Mar 19]. Available from: https://bnf.nice.org.uk/treatment-summary/beta-adrenoceptor-blocking-drugs.html

40. CPRD @ Cambridge - Code Lists [Internet]. Primary Care Unit. [cited 2018 Mar 19]. Available from: http://www.phpc.cam.ac.uk/pcu/cprd_cam/codelists/

41. National Institute for Health and Care Excellence. Hypertension in adults: diagnosis and management. [Internet]. 2011. Available from: http://www.nice.org.uk/guidance/cg127/resources/hypertension-in-adults-diagnosis-and-management-35109454941637

42. Kehoe PG, Blair PS, Howden B, Thomas DL, Malone IB, Horwood J, et al. The Rationale and Design of the Reducing Pathology in Alzheimer’s Disease through Angiotensin TaRgeting (RADAR) Trial. J Alzheimers Dis JAD. 2018;61(2):803–14.

43. Mukund Lad. The English Indices of Deprivation 2010. 2011.

44. Charlson ME, Pompei P, Ales KL, MacKenzie CR. A new method of classifying prognostic comorbidity in longitudinal studies: development and validation. J Chronic Dis. 1987;40(5):373–83.

45. Khan NF, Perera R, Harper S, Rose PW. Adaptation and validation of the Charlson Index for Read/OXMIS coded databases. BMC Fam Pract. 2010 Jan 5;11:1.

46. Lawlor B, Segurado R, Kennelly S, Olde Rikkert MGM, Howard R, Pasquier F, et al. Nilvadipine in mild to moderate Alzheimer disease: A randomised controlled trial. PLoS Med. 2018 Sep;15(9):e1002660.

47. Barthold D, Joyce G, Wharton W, Kehoe P, Zissimopoulos J. The association of multiple anti-hypertensive medication classes with Alzheimer’s disease incidence across sex, race, and ethnicity. PLOS ONE. 2018 Nov 1;13(11):e0206705.

