## Supplement for "Comparison of antihypertensive drug classes for dementia prevention"

### Supplementary Material

**Supplementary Text 1: Interpretation of bias scatter plots**

To assess bias, we constructed bias scatter plots for each outcome, which compare the association of each covariate with the exposure (obtained from multivariable linear regression analysis) and the instrument (obtained from instrumental variable analysis). If all points on these plots were on the x-axis then this would indicate that the IV analysis would be less biased than multivariable linear regression analysis. Points above the x-axis but below the x=y line would indicate bias in both analyses that is greater in the multivariable linear regression analysis estimate, while points above the x=y line would indicate bias in both analyses that is greater in the instrumental variable analysis estimate. Points on the x=y line would indicate that the bias is equal for the two analyses.

**Supplementary Text 2: Overlap with previous CPRD studies**

Two of the existing studies also make use of the CPRD. The first, by Davies et al, investigated the effects of angiotensin converting enzyme inhibitors and angiotensin-II receptor blockers, compared with other antihypertensives, on various dementia outcomes. (1) There is a small overlap between the present study and Davies et al, which we have estimated to be 5.2% at most (48,363 new users of antihypertensives in Davies et al vs 849,378 new users of antihypertensives in the present study). The second, by Goh et al, compared the effects of angiotensin-II receptor blockers and angiotensin-converting enzyme inhibitors against each other in relation to dementia as a single outcome. (2) As they did not consider other antihypertensive drug classes as an exclusion criterion, they had a much larger sample of 426,089 participants (as opposed to 221,421 participants) exposed to angiotensin-II receptor blockers and angiotensin-converting enzyme inhibitors. This made it difficult to calculate the overlap as many of these patients are likely to have been exposed to other antihypertensives. However, we do know that there were 50,404 participants assigned to the drug classes angiotensin-II receptor blockers and angiotensin-converting enzyme inhibitors in our analysis that were not present in the Goh et al study. This is because they were prescribed after 2010, i.e. after the final data extract for the Goh et al study, so will not have been included in their analysis. Despite the potential overlap of some of the data used in the present study with these studies in the literature, the study design and analysis differ considerably between them.

#### Supplementary Table 1: Amendments to the study protocol

| | |
|---|---|
| Exposure | `Centrally acting antihypertensives' are primarily used for acute events, while `Loop diuretics' are primarily used for heart failure, and so have been excluded from the analysis. We have also combined `Potassium-sparing diuretics and aldosterone antagonists' and `Thiazides and related diuretics' into a single category titled `Diuretics' as prescriptions for the former in the data extract were rare. |
| Control | Instead of using beta-adrenoceptor blocking drugs as our reference drug class for all analyses, we have used each drug class as the reference drug class in turn and presented all of the results in a matrix |

The original study protocol is available online. (3)

#### Supplementary Table 2: Fulfilment of IV study reporting guidelines

| Item | Fulfilment |
|---|---|
| State which population target parameter the study aims to estimate (eg, local average treatment effect, effect of treatment on the treated) and the assumptions on which it depends (eg, monotonicity or no effect modification). | The following statements are made in the paper: "the results were interpreted as the effect among patients whose prescription was affected by their physicians' preference (known as the local average treatment effect)" and "Instrumental variable analysis requires that the instrument: (i) be associated with the exposure of interest; (ii) affect the outcome only through its effect on the exposure of interest; and (iii) have no common causes with the outcome. To obtain a point estimate for this analysis, we also make a fourth assumption of monotonicity." |
| Report the association of instruments and exposure using a partial F-statistic. | The Cragg-Donald F statistic has been presented alongside the results for each of our analyses. |
| Report and test the association of observed potential confounding factors with both the exposure and the instrument. | See section `Assessment of bias' in the paper. |
| With multiple instruments report the test for overidentifying restrictions, ie, the Sargan or the Hansen test. | This is not applicable to this study as we use single instruments. |
| For binary outcomes, exposures, and instruments, report a tabulation of the frequencies of each combination of instrument, exposure, and outcome, so readers can reconstruct basic results. | This is not applicable to this analysis as we have a categorical instrument and would need to report 32 instrument-exposure-outcome combinations per analysis conducted. |
| When using generalized linear models with binary outcomes, always use robust (sandwich estimators) or bootstrapped standard errors and take clustering of study participants into account where necessary. | The analysis used Stata's ivreg2 command with 'robust' specified and clustering according to the physicians' staff ID. The analysis was conducted in Stata version 15MP. |

#### Supplementary Table 3: Covariates adjusted for in the multivariable logistic regression analysis

| Covariate | How was the covariate defined? |
|---|---|
| Previous history of coronary heart disease | Presence of one or more relevant Read codes on record. |
| Previous history of coronary-bypass surgery | Presence of one or more relevant Read codes on record. |
| Previous history of cerebrovascular disease (including stroke) | Presence of one or more relevant Read codes on record. |
| Chronic illness, including cancer and arthritis | Charlson index implemented using Read code lists. (4,5) Code lists based on those by Taylor et al. (6) |
| Socioeconomic position | 2010 English Index of Multiple Deprivation (IMD) at the 'twentile' level, where 1 represents the least deprived and 20 the most deprived. |
| Consultation rate | Calculated by dividing the total number of clinic visits by the length of the patient record prior to the index date to give an average annual rate. |
| Alcohol status | Recorded value (current, former or never). |
| Smoking status | Most recent of recorded value (current, former or never) or Read code indicating a recorded value. Code lists based on those by Wright et al. (7) |
| BMI | Recorded value if available, or a calculated value using the last recorded height and weight measurements. Measurements taken before the age of 25 were excluded to ensure adult measurements were used. |

**Supplementary Table 4: Exposure by age and sex**

| Drug class / Age group | 40-49 | 50-59 | 60-69 | 70-79 | 80-89 | 90-99 | 100+ | Total |
|---|---|---|---|---|---|---|---|---|
| Alpha-adrenoceptor blockers | 5,979<br>(93.4) | 15,289<br>(97.1) | 22,482<br>(98.0) | 16,141<br>(97.5) | 6,653<br>(96.3) | 806<br>(94.7) | 10<br>(100.0) | 67,360<br>(97.0) |
| Angiotensin-II receptor blockers | 42,697<br>(62.1) | 57,932<br>(62.9) | 45,726<br>(60.0) | 32,124<br>(50.4) | 14,881<br>(42.1) | 2,482<br>(32.6) | 49<br>(20.4) | 195,891<br>(58.0) |
| Angiotensin converting enzyme inhibitors | 2,994<br>(63.1) | 4,366<br>(61.5) | 3,742<br>(55.2) | 2,479<br>(44.7) | 1,010<br>(36.2) | 125<br>(21.6) | 1<br>(0.0) | 14,717<br>(55.3) |
| Beta-adrenoceptor blockers | 77,343<br>(39.3) | 69,672<br>(45.7) | 49,775<br>(48.2) | 30,948<br>(43.0) | 11,579<br>(35.7) | 1,522<br>(26.2) | 25<br>(20.0) | 240,864<br>(43.2) |
| Calcium channel blockers | 16,569<br>(50.6) | 31,237<br>(55.8) | 45,022<br>(53.8) | 32,318<br>(42.7) | 12,832<br>(34.9) | 1,721<br>(23.7) | 31<br>(16.1) | 139,730<br>(49.2) |
| Diuretics | 22,189<br>(33.3) | 38,671<br>(39.2) | 47,552<br>(40.9) | 45,820<br>(34.2) | 22,961<br>(28.9) | 3,660<br>(23.1) | 93<br>(11.8) | 180,946<br>(36.0) |
| Vasodilator antihypertensives | 2,440<br>(99.2) | 3,381<br>(99.5) | 3,033<br>(99.7) | 907<br>(98.6) | 103<br>(91.3) | 5<br>(40.0) | 1<br>(0.0) | 9,870<br>(99.3) |
| Total | 170,211<br>(48.5) | 220,548<br>(55.2) | 217,332<br>(56.3) | 160,737<br>(47.7) | 70,019<br>(40.5) | 10,321<br>(31.5) | 210<br>(19.5) | 849,378<br>(51.2) |

Each cell contains the total number of patients prescribed the drug class of interest in that age group. The number in brackets states the percentage of that number that are male.

**Supplementary Table 5: Results from all analyses**

Due to the size of this table, it is not included here but is available from GitHub via the following link: https://github.com/venexia/repurposing-antihypertensives-dementia

**Supplementary Figure 1: Study design diagram**

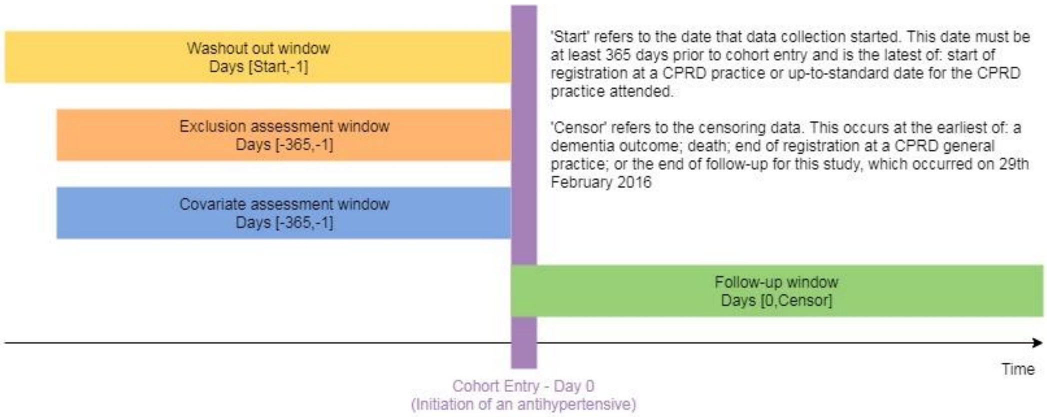

**Supplementary Figure 2: Decision tree for outcome definitions**

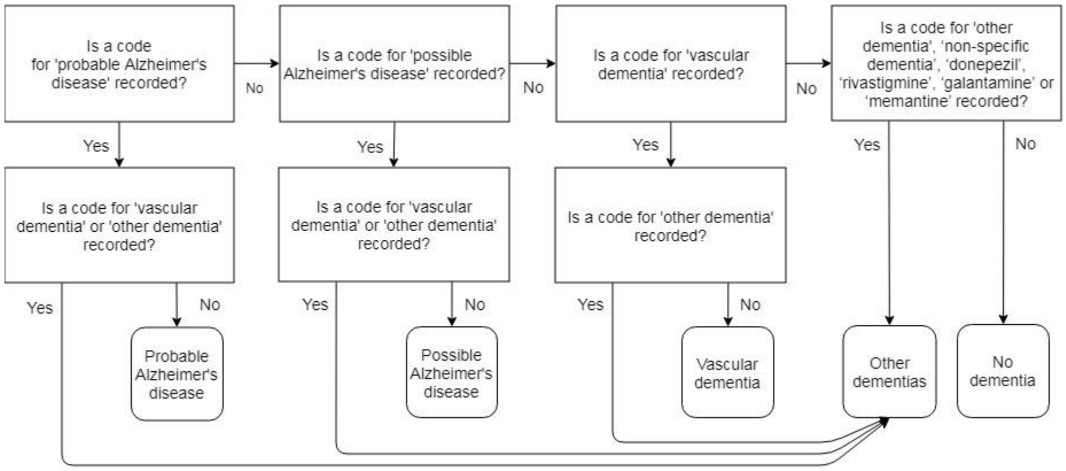

**Supplementary Figure 3: Attrition of patients in the analysis cohort**

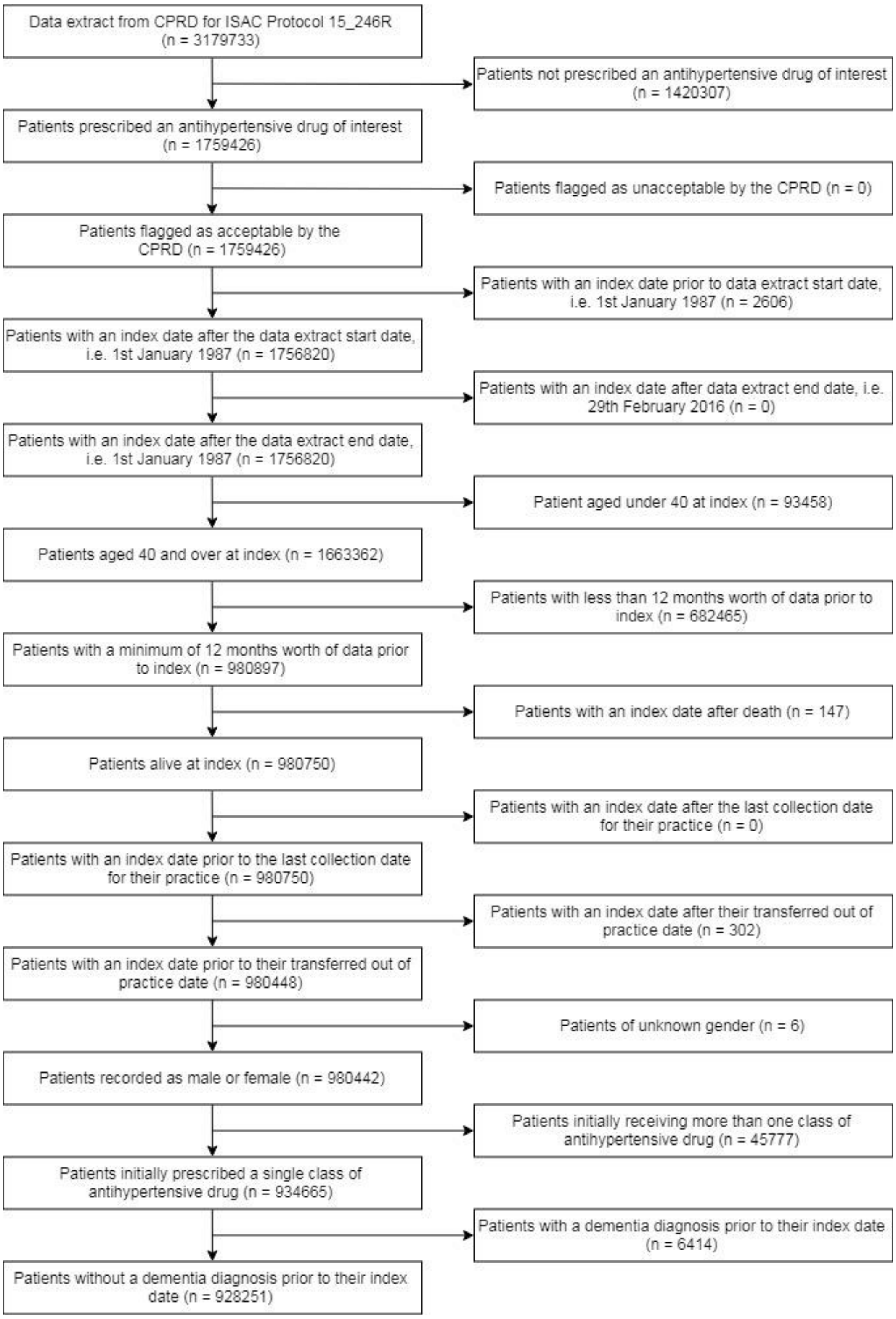

**Supplementary Figure 4: Multivariable logistic regression results**

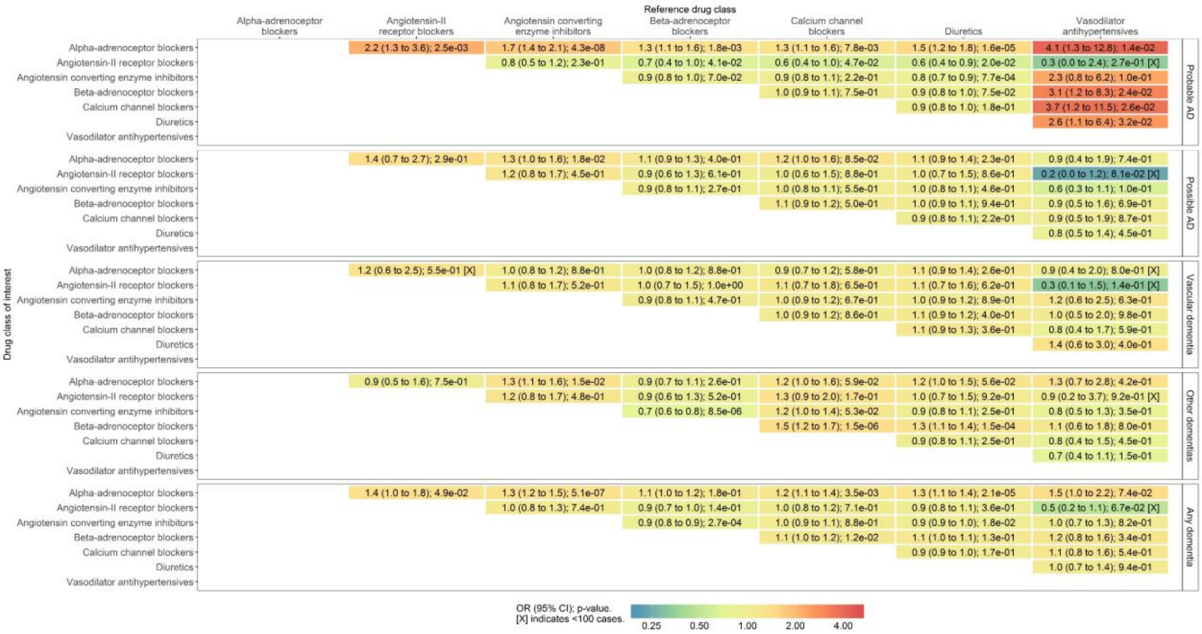

For a drug class of interest, i.e. a ‘row’, estimates below zero indicate a protective effect of that drug class. For a reference drug class, i.e. a ‘column’, estimates above zero indicate a protective effect of that drug class. This matrix is antisymmetric, so the values not presented are the negation of those that are presented.

**Supplementary Figure 5: Bias scatter plot for covariates in the any dementia analysis**

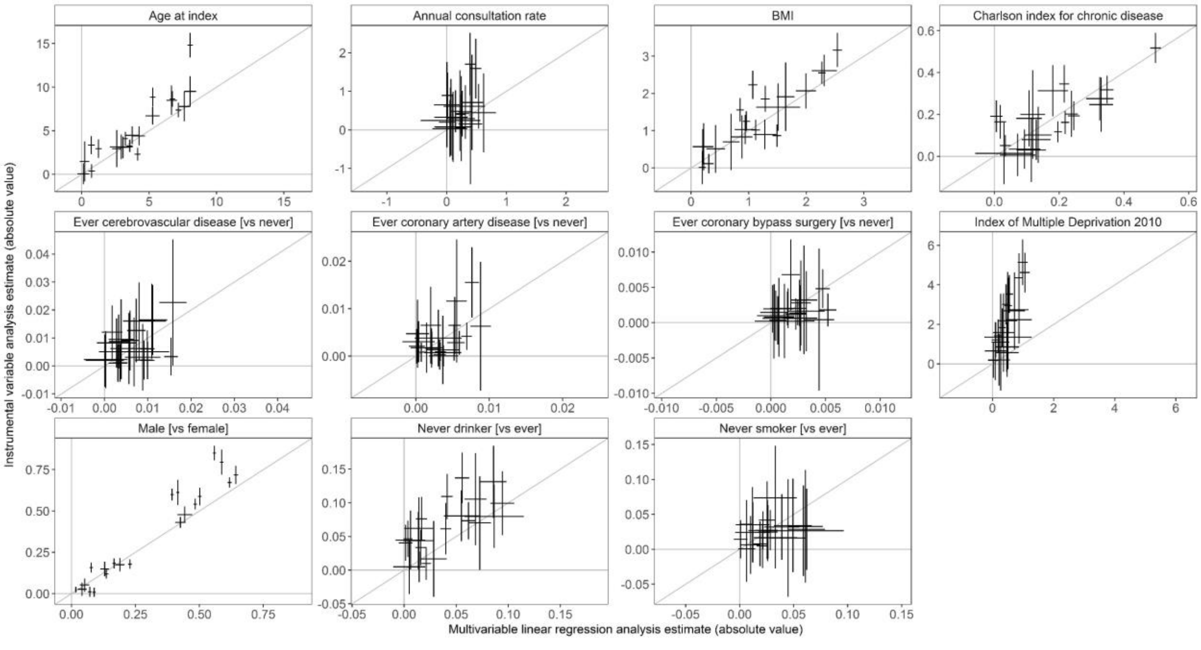

Each point on a scatter plot represents an individual analysis with the outcome 'any dementia'. This plot is representative of the results obtained for all outcomes.

**Supplementary Figure 6: IV estimates after adjustment for socioeconomic position**

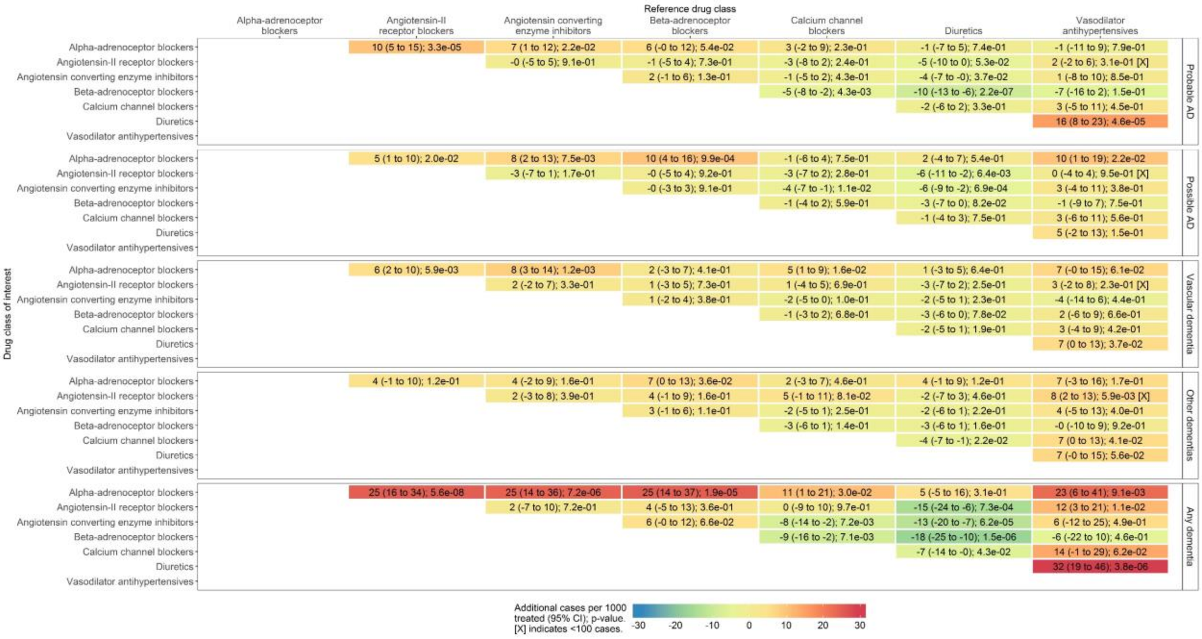

F greater than 2530 for all analyses. For a drug class of interest, i.e. a ‘row’, negative estimates indicate a protective effect of that drug class. For a reference drug class, i.e. a ‘column’, positive estimates indicate a protective effect of that drug class. This matrix is antisymmetric, so the values not presented are the negation of those that are presented.

**Supplementary Figure 7: IV estimates after adjustment for BMI**

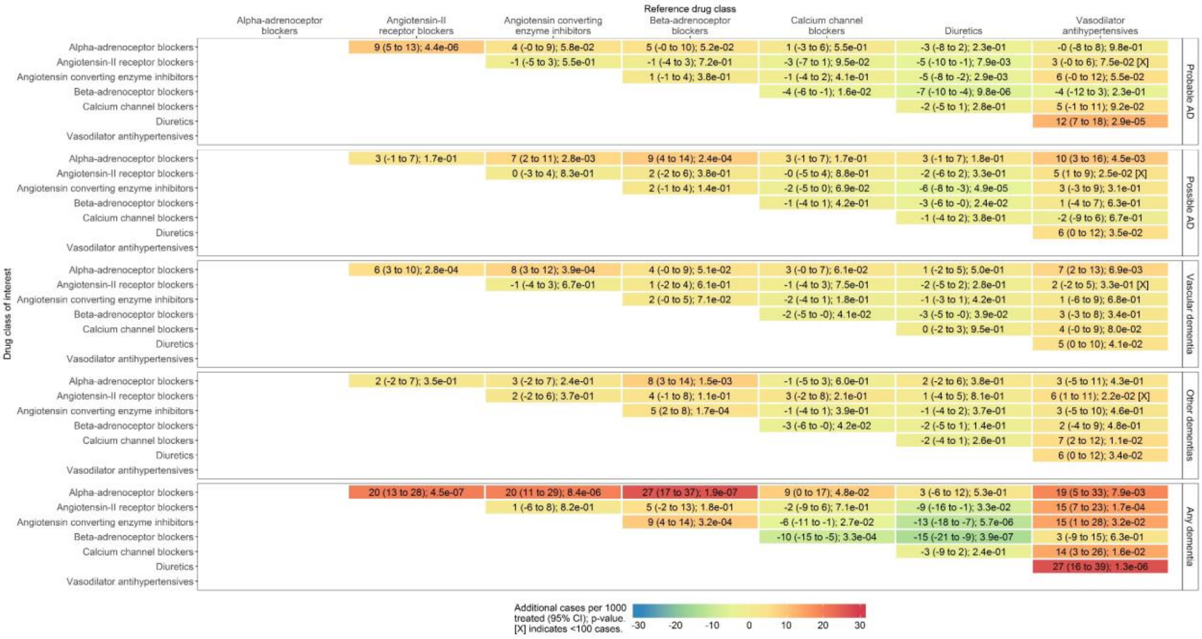

F greater than 3890 for all analyses. For a drug class of interest, i.e. a ‘row’, negative estimates indicate a protective effect of that drug class. For a reference drug class, i.e. a ‘column’, positive estimates indicate a protective effect of that drug class. This matrix is antisymmetric, so the values not presented are the negation of those that are presented.

**Supplementary Figure 8: IV estimates after adjustment for chronic disease**

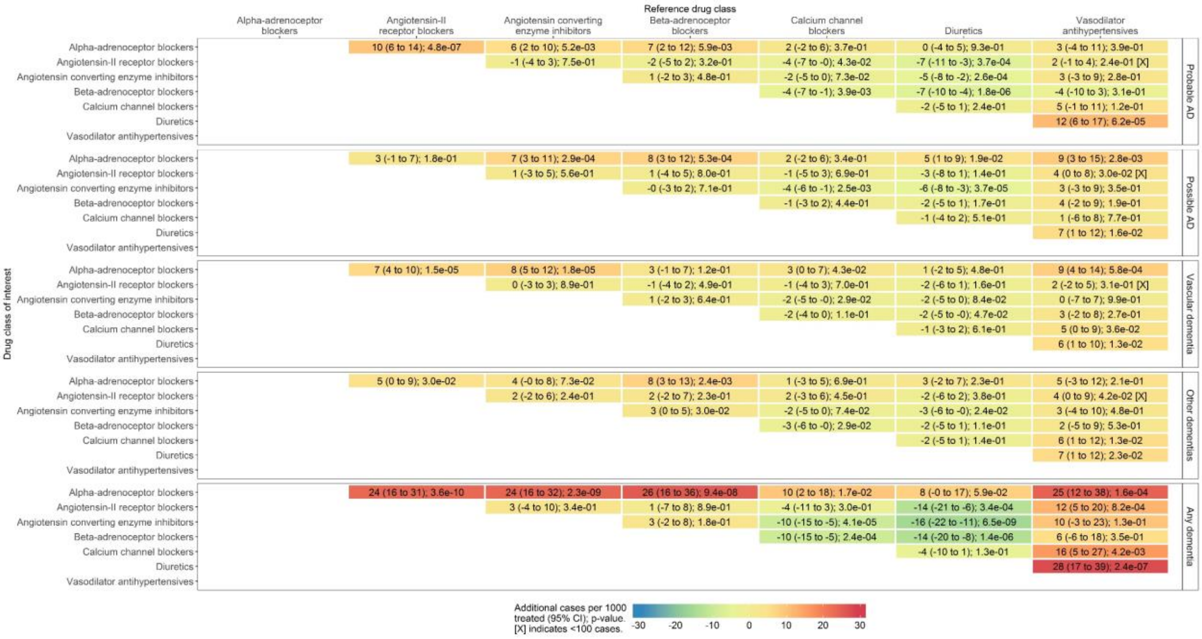

F greater than 4499 for all analyses. For a drug class of interest, i.e. a ‘row’, negative estimates indicate a protective effect of that drug class. For a reference drug class, i.e. a ‘column’, positive estimates indicate a protective effect of that drug class. This matrix is antisymmetric, so the values not presented are the negation of those that are presented.

**Supplementary Figure 9: IV estimates after adjustment for sex**

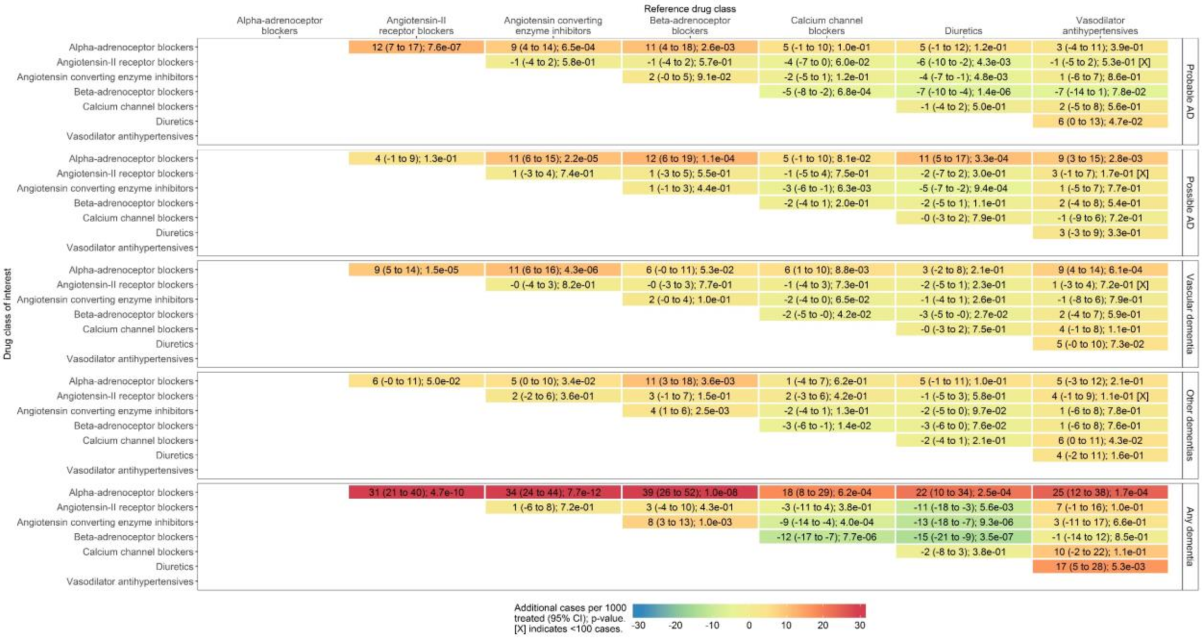

F greater than 4416 for all analyses. For a drug class of interest, i.e. a ‘row’, negative estimates indicate a protective effect of that drug class. For a reference drug class, i.e. a ‘column’, positive estimates indicate a protective effect of that drug class. This matrix is antisymmetric, so the values not presented are the negation of those that are presented.

**Supplementary Figure 10: IV estimates after adjustment for age**

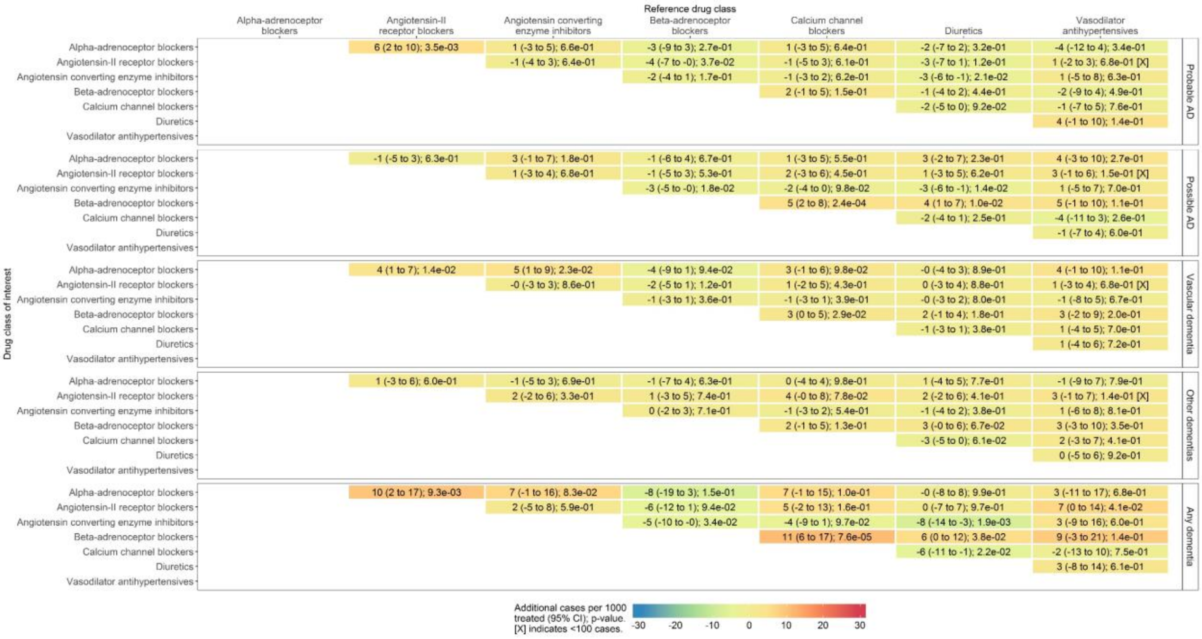

F greater than 4094 for all analyses. For a drug class of interest, i.e. a ‘row’, negative estimates indicate a protective effect of that drug class. For a reference drug class, i.e. a ‘column’, positive estimates indicate a protective effect of that drug class. This matrix is antisymmetric, so the values not presented are the negation of those that are presented.

**Supplementary Figure 11: IV estimates without patients diagnosed with anxiety in the same consultation**

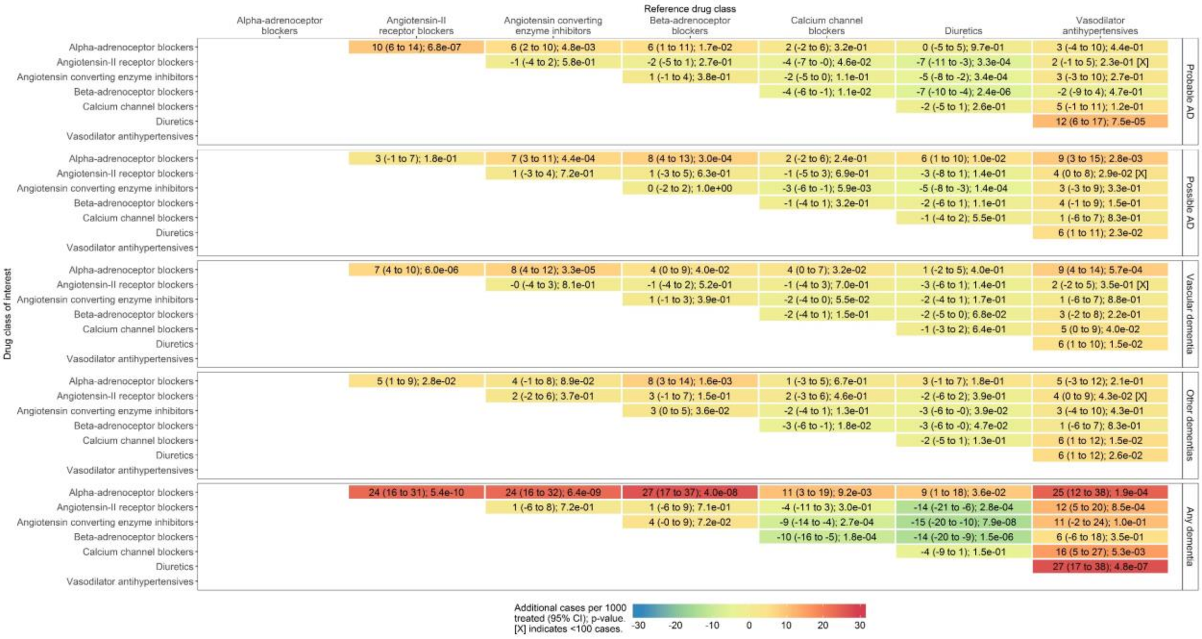

F greater than 4705 for all analyses. For a drug class of interest, i.e. a ‘row’, negative estimates indicate a protective effect of that drug class. For a reference drug class, i.e. a ‘column’, positive estimates indicate a protective effect of that drug class. This matrix is antisymmetric, so the values not presented are the negation of those that are presented.

### Supplementary Figure 12: IV estimates without patients who had a low dose initial prescription

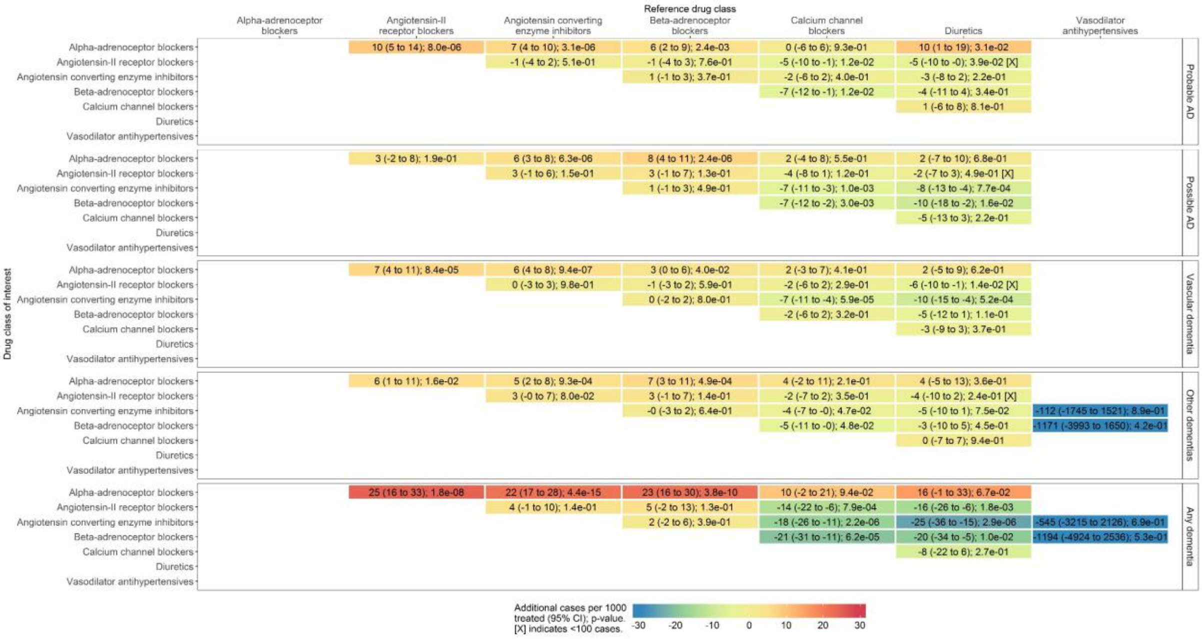

F greater than 13 for all analyses. Missing cells due to insufficient sample size to run the analysis. For a drug class of interest, i.e. a ‘row’, negative estimates indicate a protective effect of that drug class. For a reference drug class, i.e. a ‘column’, positive estimates indicate a protective effect of that drug class. This matrix is antisymmetric, so the values not presented are the negation of those that are presented.

### Supplementary Figure 13: IV estimates for patients aged 55 and over at index

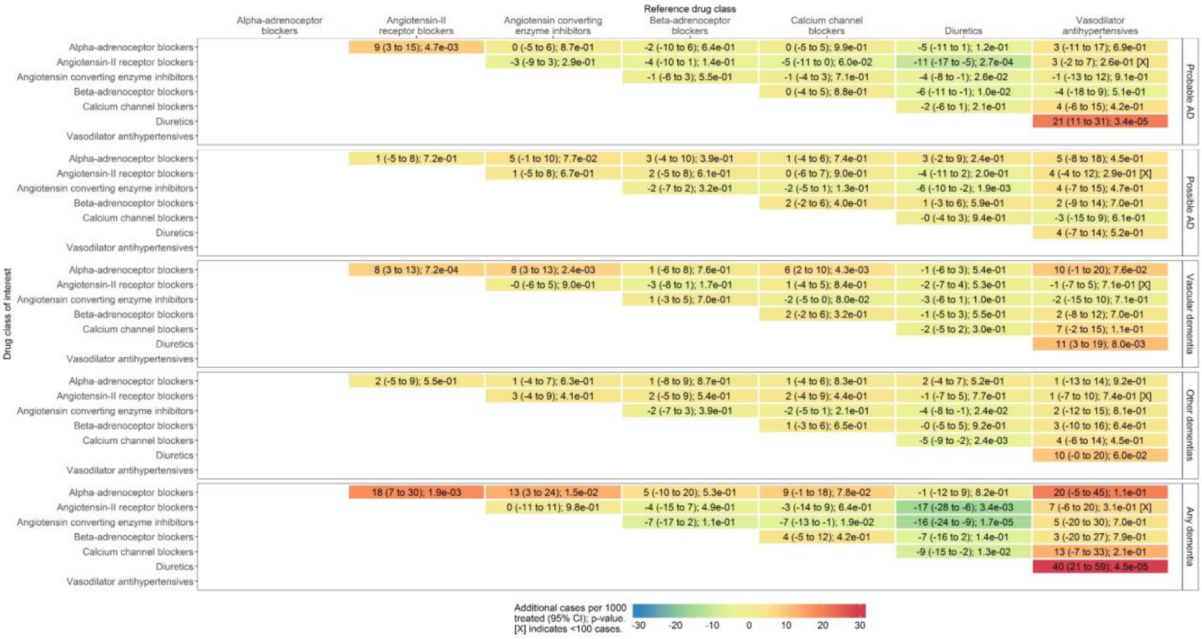

F greater than 1956 for all analyses. For a drug class of interest, i.e. a ‘row’, negative estimates indicate a protective effect of that drug class. For a reference drug class, i.e. a ‘column’, positive estimates indicate a protective effect of that drug class. This matrix is antisymmetric, so the values not presented are the negation of those that are presented.

### References

1. Davies NM, Kehoe PG, Ben-Shlomo Y, Martin RM. Associations of anti-hypertensive treatments with Alzheimer's disease, vascular dementia, and other dementias. J Alzheimers Dis. 2011;26(4):699.

2. Goh KL, Bhaskaran K, Minassian C, Evans SJW, Smeeth L, Douglas IJ. Angiotensin receptor blockers and risk of dementia: cohort study in UK Clinical Practice Research Datalink. Br J Clin Pharmacol. 2015 Feb 1;79(2):337–50.

3. Walker VM, Davies NM, Jones T, Kehoe PG, Martin RM. Can commonly prescribed drugs be repurposed for the prevention or treatment of Alzheimer's and other neurodegenerative diseases? Protocol for an observational cohort study in the UK Clinical Practice Research Datalink. BMJ Open. 2016 Dec 1;6(12):e012044.

4. Charlson ME, Pompei P, Ales KL, MacKenzie CR. A new method of classifying prognostic comorbidity in longitudinal studies: development and validation. J Chronic Dis. 1987;40(5):373–83.

5. Khan NF, Perera R, Harper S, Rose PW. Adaptation and validation of the Charlson Index for Read/OXMIS coded databases. BMC Fam Pract. 2010 Jan 5;11:1.

6. Taylor GM, Taylor AE, Thomas KH, Jones T, Martin RM, Munafò MR, et al. The effectiveness of varenicline versus nicotine replacement therapy on long-term smoking cessation in primary care: a prospective cohort study of electronic medical records. Int J Epidemiol. 2017 Dec 1;46(6):1948–57.

7. Wright AK, Kontopantelis E, Emsley R, Buchan I, Sattar N, Rutter MK, et al. Life Expectancy and Cause-Specific Mortality in Type 2 Diabetes: A Population-Based Cohort Study Quantifying Relationships in Ethnic Subgroups. Diabetes Care. 2017 Mar 1;40(3):338–45.
